# Contact, travel, and transmission: The impact of winter holidays on influenza dynamics in the United States

**DOI:** 10.1101/055871

**Authors:** Anne Ewing, Elizabeth C. Lee, Cécile Viboud, Shweta Bansal

## Abstract

**Background:** The seasonality of influenza is thought to vary according to environmental factors and human behavior. During winter holidays, potential disease-causing contact and travel deviate from typical patterns, and we aim to understand these changes on age-specific and spatial flu transmission.

**Methods:** We characterized the changes to transmission and epidemic trajectories among children and adults in a spatial context before, during, and after the winter holidays among aggregated physician medical claims in the United States from 2001 to 2009 and among synthetic data simulated from a deterministic, age-specific spatial metapopulation model.

**Results:** Winter holidays reduced flu transmission and delayed the trajectory of flu season epidemics. The holiday period itself observed a shift in relative risk of disease from children towards adults. Model results indicated that holidays delay epidemic peaks and synchronize incidence across locations, and contact reductions from school closures rather than age-specific mixing and travel produce these observed holiday dynamics.

**Conclusions:** Winter holidays delay seasonal influenza epidemic peaks due to changes in contact patterns. These findings may improve the future design of influenza intervention strategies, such as the proper timing and duration of school closures, and the spatial and demographic allocation of vaccines.

## Introduction

Influenza epidemics are characterized by large variation in disease burden across seasons and across locations within a given season [1]. While we do not fully understand what drives this variation, contact and travel patterns have been observed to influence local and global influenza transmission [2, 3, 4, 5]. Winter holidays alter typical contact and travel patterns through school closures and holiday travel and occur early in typical flu seasons in temperate climates, yet the measurement and the subsequent effects of these temporary behavioral changes on seasonal flu remain poorly understood.

Consideration of age group patterns is an important component to understanding the transmission and relative disease burden of influenza. Empirical contact surveys have illustrated that individuals tend most to associate with others in a similar age group and school-aged children tend to have the greatest number of potential disease-causing contacts [6, 7]. Additionally, modeling studies have demonstrated that social mixing by age is sufficient to capture much of the heterogeneity in contact across populations [8]. In large population settings, school-aged children are thought to drive local transmission of flu due to their large number of contacts, while adults are thought to seed flu in different locations due to their global mobility [3, 4, 9].

Since schools and school-aged children are of particular importance for influenza transmission, temporary school closures are commonly considered as a reactive intervention in pandemic and severe flu seasons [10, 11]. However, empirical studies of the effect of these interventions varies from no significant effect on flu transmission, to reducing transmission by 29% in children alone [12, 13, 14]. Due to these mixed results, the impact on children and the subsequent trickle-down effects to other age groups remains unclear. While school holidays have similarities to closures, they occur at predetermined times and induce changes to both contact and travel patterns. In the United States, the Christmas holiday occurs in late December and U.S. epidemics typically start in December and peak in February; changes in flu transmission during the winter holidays could crucially affect the resulting flu epidemic [15]. The number of contacts between children decreases as children are out of school for the holiday [16], and mixed results suggest that winter school holidays reduce or delay the risk of influenza among school-aged children by 33–42% [17, 18] and periods around the holidays experience high variability in ILI across seasons [19].

Local and global travel are mechanisms by which respiratory pathogens are commonly thought to spread, and travel patterns have long been studied to understand the spatial spread of diseases. Winter holidays are characterized by increases in visits to friends and family that cause deviations from typical travel patterns, and in the U.S., there are notable increases in travel among children and in the volume of long-distance travel [20, 21]. Human movement is tied to the phylogeographic spread of influenza viruses [22], and substantial evidence suggests that travel can influence the spatial spread and timing of influenza epidemics [3, 23, 24, 25, 4, 26]. However, travel restrictions have been shown to produce little to no effect on the spread of pandemic influenza [27, 28, 29]

Our work aims to determine the impact of changing contact and travel patterns during winter holidays on influenza transmission and the resulting epidemic trajectories. With U.S. medical claims data, we examine changes in flu transmission during and after the holidays, and characterize common patterns in the rates of influenza-like illness among school-aged children and working-aged adults during the holiday period across multiple flu seasons. To understand the mechanisms behind these empirical patterns, we create a parsimonious age-specific spatial metapopulation model to study the interactions among child and adult populations and the importance of human travel in spatial spread. Using the model, we examine two hypotheses that may work independently or in concert: 1) holiday changes to travel patterns temporarily synchronize epidemics across locations (subpopulation dynamics) [30, 31], or 2) holiday reductions in contact patterns damp transmission to such low levels that all epidemics ‘reset’ to the beginning of their trajectories, thus creating an appearance of synchrony [18, 17]. Using a combination of empirical and theoretical approaches, our work highlights the significant role of the holidays on shaping influenza seasonal dynamics, and has implications for influenza control through vaccination prioritization and school closures.

## Methods

We used reports of influenza-like illness (ILI) to explore spatial and age-specific patterns of seasonal influenza activity around the winter holidays in the United States. Motivated by these empirical observations, we then tested hypotheses about the mechanisms that may be driving holiday flu dynamics through a simple epidemiological model. Here, we introduce the empirical data, mathematical model structure, and the measures through which we characterized spatial and age-specific epidemiological patterns with the empirical and simulated data. (More details are available in the SM sections S1 and S2).

### Medical claims data

Weekly visits for ILI and any diagnosis from October 2001 to May 2009 were obtained from a records-level database of CMS-1500 US medical claims managed by IMS Health and aggregated to three-digit U.S. zipcode prefixes (zip3s), where ILI was derived from a set of International Classification of Diseases, Ninth Revision (ICD-9) codes and validated at multiple geographic scales, as described elsewhere [32]. To account for temporal variation in health care-seeking (e.g., doctor’s office closures, lower probability of seeking care for illness during holidays), weekly ILI visits were divided by weekly all-cause patient visits (i.e. a visit is included regardless of its purpose) in the medical claims database and standardized by population size to calculate an *ILI incidence ratio* (See SM section S1.1, Figure S1) [32]. For age-specific analyses, we calculate ILI incidence ratios for individuals 5–19 years old to represent children and for individuals 20–69 years old to represent adults. The epidemic period was defined as October through March, and ‘before’, ‘during’ and ‘after’ periods of the holiday were defined as two week periods, with the "during" period beginning at the Christmas holiday.

To understand the effect of the winter holidays on flu transmission in the empirical data, we estimated the effective reproductive number (*R*_*t*_), the average number of secondary cases generated by each infected individual under the conditions at time *t*, over weekly periods during the eight flu seasons from 2001–2002 through 2008–2009. (Details on the calculation of *R*_*t*_ can be found in SM section S1.3).

### Metapopulation model

We used an epidemiological model to simulate flu epidemics with and without holiday-associated behavioral changes to contact and travel patterns. Our model was adapted from an age-specific metapopulation model that 1) incorporates contact between children and adults, and 2) is spatially divided into metropolitan areas linked through air traffic flows [4, 33]. Infection followed an SIR (Susceptible, Infected, Recovered) disease progression, and the entire population was assumed to be susceptible at the start of an outbreak. Each model run was seeded with one child infection in a single spatial area. Disease spread deterministically and in discrete time steps according to age-specific contact patterns, and infection reached additional metro areas through travel. All model results shown are an average across all possible seeds. (Details on the demographic and travel data used to parameterize the model can be found in SM section S2.)

### Experimental design

To mimic holiday-associated behaviors in the model, we altered age-specific contact and travel parameters during a predetermined 14-day holiday period (See SM section 2.3). We identified the holiday period based on the average number of days between Christmas and the epidemic peak in the national empirical data, and the ‘before’, ‘during’ and ‘after’ periods of the holiday were defined as two week periods, with the "during" period beginning at the holiday. We also carried out sensitivity analyses to compare epidemiological patterns when holiday period contact rates and timing were altered (See SM sections S3.2 and S3.3). In the *school closure model*, we altered each value of the baseline contact matrix according to empirical survey data that reported age-specific contact rates during school holidays. Specifically, during the holiday period, the total contact rate was reduced in both age groups and the rate of child to adult contact increased proportional to the total number of child contacts [16]. In the *travel model*, during the holiday period, we altered the travel-based connectivity between metropolitan areas based on air traffic patterns from December 2005, and by increasing the fraction of child travelers, r, to 15% [20]. Generally speaking, holiday travel observed a greater volume of travelers pushed through fewer locations than baseline travel (See SM section S2.4). The *holiday model* combined the changes associated with both the school closure and travel models.

## Results

We first explore empirical patterns of influenza burden and transmission during and after the holiday period in the United States across multiple flu seasons. We then consider results from an age-specific spatial metapopulation model to systematically understand the impact of holiday-related school closures and changes to travel patterns on the spatial and age-specific spread of influenza.

### Characterizing empirical influenza patterns during the holidays

Here, we report age-specific and spatial patterns of influenza transmission based on U.S. medical claims data for influenza-like illness (ILI) during the months around the Christmas holiday, after having corrected for variation in reporting rates. (See Methods for details.)

During the weeks following Christmas, we observed temporary reductions in U.S. flu activity across eight seasons, even after accounting for holiday-associated reductions in healthcare-seeking (Figures 1A and S1). Flu transmission, as measured by the mean effective reproductive number (*R*_*t*_) (see Methods for further details), decreased by approximately 15% (reduction from *R*_*t*_ ~ 1.1 to ~ 0.9) in most seasons and fell below the epidemic threshold (*R*_*t*_ = 1) immediately following Christmas. Within a few weeks, flu transmission exceeded the epidemic threshold and rebounded to pre-holiday levels (Figure 1B). These patterns were observed consistently within our study period, including the notably early 2003–04 flu season. This pattern was also observed across smaller spatial regions, specifically across zipcodes sharing the first three digits (zip3) in the United States (Figure S2). Analysis of viral surveillance data on influenza-positive laboratory confirmations verified that influenza was indeed circulating around the Christmas holiday for each season in our study period, so decreases in ILI and *R*_*t*_ may be attributed to influenza dynamics; similarly, we did not investigate Thanksgiving holiday dynamics because influenza circulation was quite limited during this period (Figure S3).

**Figure 1:**
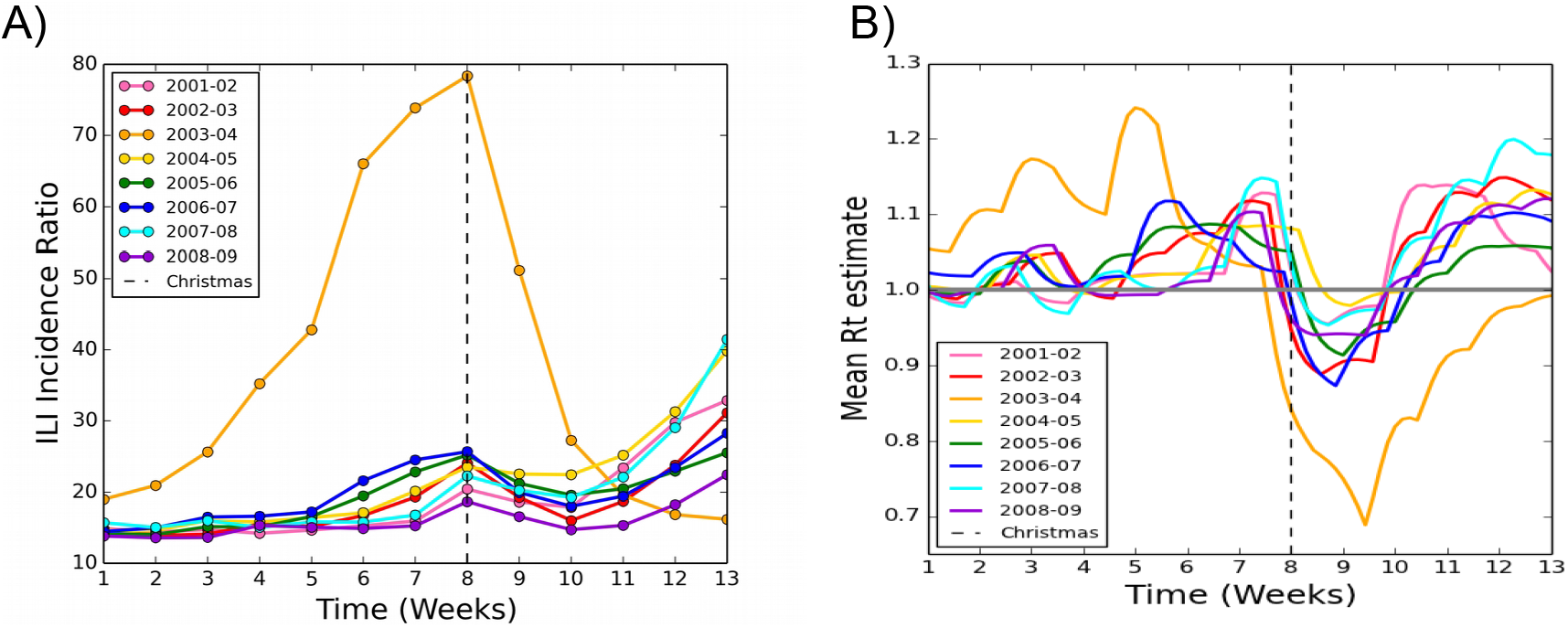
Decreases in transmission are observed following Christmas. **A**) National ILI incidence ratio (ILI cases per total visits per 100,000 population) calculated using weekly ILI medical claims data over time in weeks from the first week in November to the last week in January for flu seasons from 2001 to 2009. The week of Christmas is marked with the vertical dotted line. **B**) National daily effective reproductive number (*R*_*t*_) over time from November to January for flu seasons from 2001 to 2009. *R*_*t*_ was calculated over seven-day windows using ILI medical claims data adjusted for health care facility closures and for care-seeking. The date of Christmas is marked with the dashed line.

We also examined weekly ILI medical claims for school-aged children (5–19 years of age) and adults (20–69 years of age) from November through January. Across the eight flu seasons, both children and adults experienced temporary declines and recoveries around the Christmas holiday (Figure 2A). However, the changes in incidence patterns were not synchronous in the two age groups, with adults experiencing a reduction only after the holiday. To examine the differential impact on the two age groups, we examined relative risk of ILI activity between children and adults. We observed that risk of disease was shifted towards adults during and after the holiday, and that these dynamics coincided with the temporary reductions in flu activity and flu transmission (Figure 2B). We posit that these patterns may be driven by altered interaction patterns due to children being home from school or families (rather than business travelers) traveling for the holidays.

**Figure 2:**
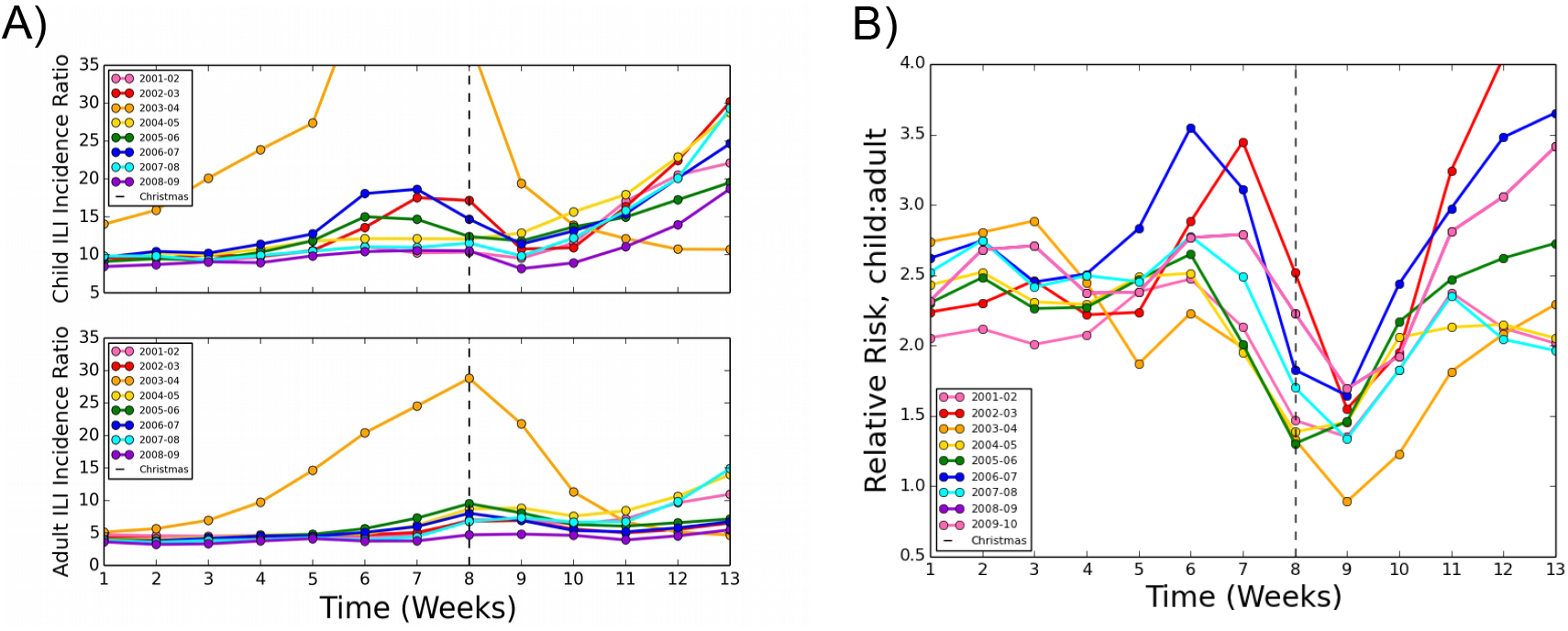
The impact of the holidays varies by age group. **A**) Age-specific ILI incidence ratio calculated from weekly ILI medical claims data from November to January for flu seasons from 2001 to 2009 among school children and adults. The week of Christmas is denoted by the vertical dotted line. **B**) ILI incidence ratio relative risk between school children and adults calculated over time in weeks from November to January medical claims data for flu seasons 2001–2009. The week of Christmas is denoted by the dashed line. A relative risk greater than 1 indicates a greater risk toward children, less than 1 indicating greater risk toward adults.

To investigate the spatial patterns of influenza spread during the holiday period, we characterized the peak timing and synchrony of ILI reports across zip3s in the medical claims data. We observed that the timing and variation of seasonal influenza peaks across zip3s in the United States is comparable for most seasons, occurring an average of five weeks after the holiday period; the early 2003–04 is an exception with the holiday period occurring after the epidemic peak in a majority of locations (Figure 3A). Additionally, the distribution of incidence across zip3 areas showed little variation from the periods before to after the holidays (Figure 3B). We hypothesize that these patterns may be driven by increased travel that homogenizes and synchronizes influenza risk across the country.

**Figure 3:**
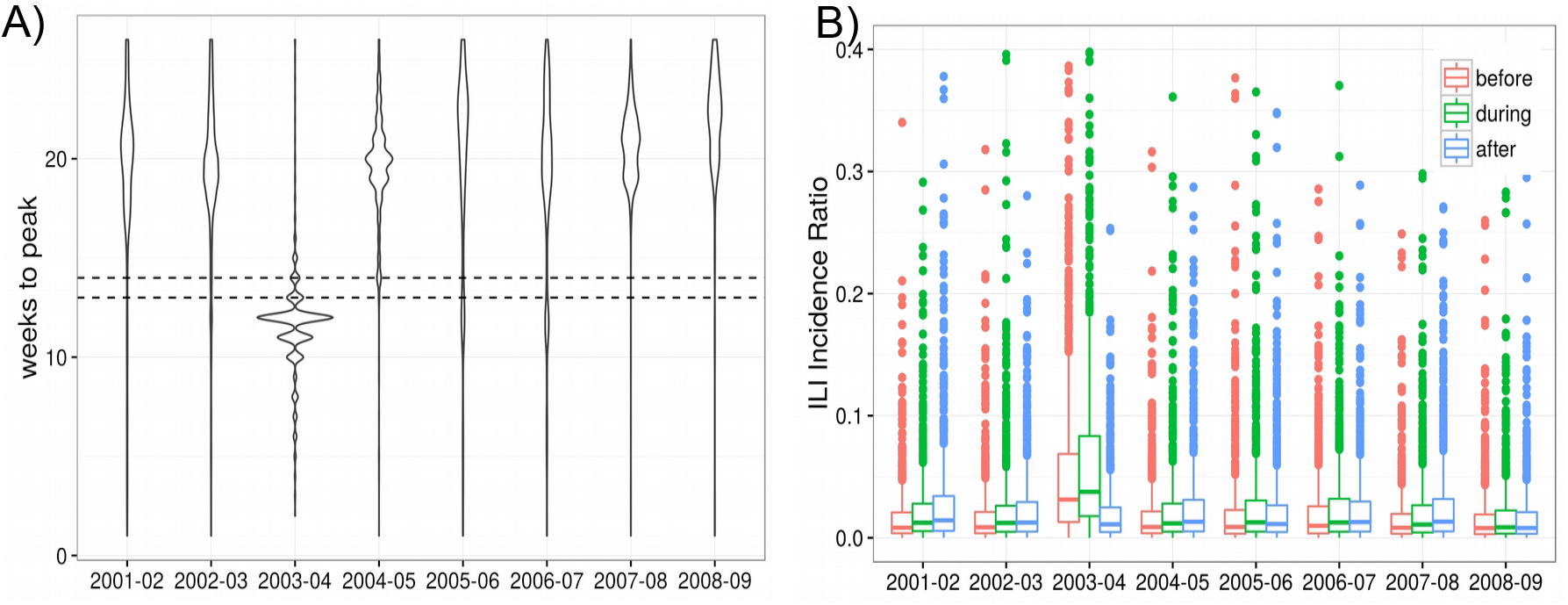
Peak timing and spatial synchrony in empirical data. **A**) Distribution of weeks to peak across all zip3s, where the flu period endures from October to March. Distributions are compared across eight flu seasons in the study period. The horizontal dashed lines highlight the holiday period. **B**) Distributions of ILI reports across all zip3s averaged for the two week durations defined as ‘before’, ‘during’, and ‘after’ the holiday periods. Distributions are compared across eight flu seasons in the study period. A small number of outlying data points are not depicted here.

### Exploring the mechanisms behind holiday flu dynamics

Informed by the empirical patterns observed in our data, we sought to identify the impact of specific behavior changes expected to occur during the holiday period. We hypothesized that two potential mechanisms influenced the age-specific and spatial patterns of influenza spread around the holidays: school closures and travel. As these mechanisms both lead to changes in individual-level and large-scale contact patterns during the holidays, we studied their impact on holiday flu dynamics in a controlled manner through the use of a mathematical model. We compared the results from three models with holiday-associated behavioral changes to a baseline model, where no behavioral changes were implemented. The holiday models implemented behavioral changes as follows: 1) increased fraction of child travelers and changes to travel volume and connectivity, according to data on altered holiday travel patterns (“travel”), 2) reduced overall number of potential disease-causing contacts for children and adults, and a greater proportion of child to adult mixing, corresponding to a holiday school closure informed by data (“school closure”), and 3) both of the above scenarios together (“holiday”). (See Methods and SM section S2 for details).

The baseline model results followed an epidemic trajectory with a single peak, while the holiday model epidemics had similar dynamics to the empirical data, illustrating a temporary drop and recovery of flu activity during and after the holiday period with a larger epidemic peak that followed (Figure 4A). The travel model produced results similar to the baseline model, while the school closure model results is largely comparable to the holiday model. As with the empirical data, children always had greater risk of flu than adults, and the risk shifted towards adults with the school closure and holiday models (Figure 4B). No such shift was observed for the baseline and travel simulations. Despite the differences in dynamics, the total epidemic sizes were comparable for all four models (Table S3).

**Figure 4:**
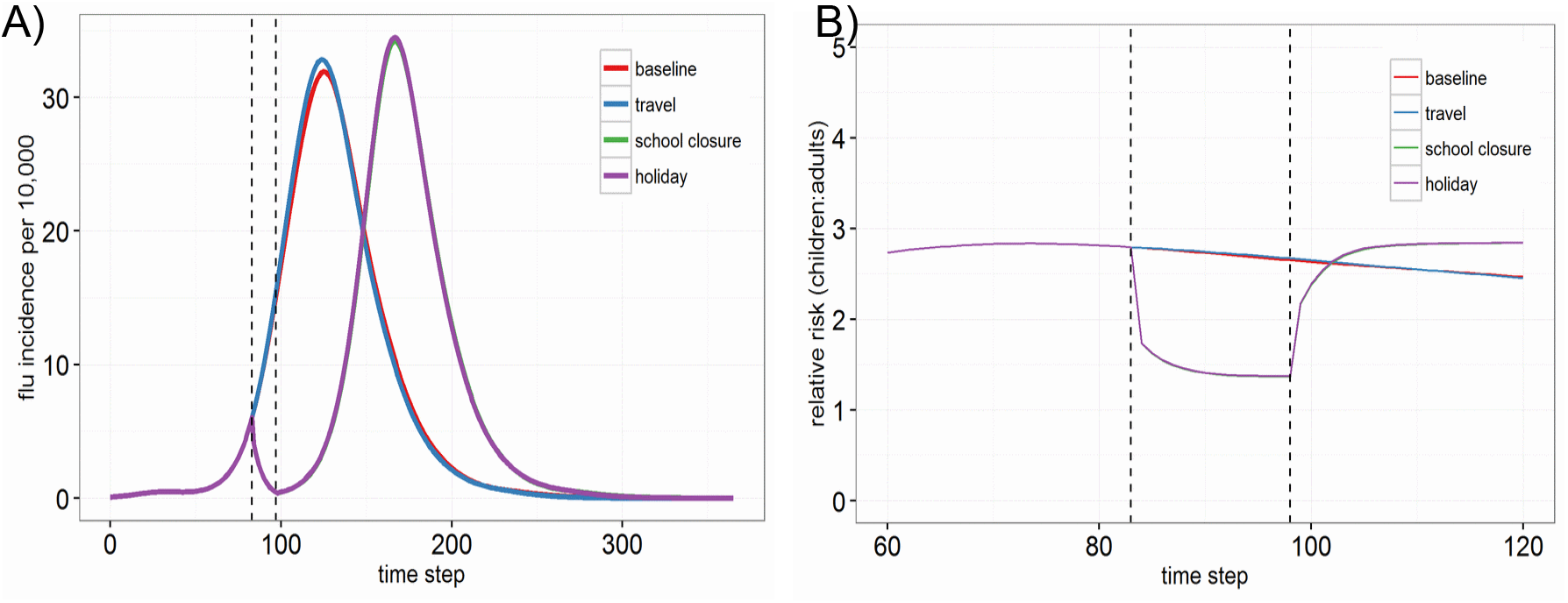
Changes to contact patterns appear to drive holiday-associated dynamics in model simulations. **A**) Total flu incidence per 10,000 population over time, averaged across all model runs. **B**) Relative risk of disease from children to adults across all locations, averaged across all model runs. Epidemic trajectories for the baseline (no changes during the holiday period), travel only, school closure only, and full holiday (travel and school closure changes) interventions are compared, and the holiday period is demarcated by the dashed black lines. The purple line shadows the green line in both figures.

To characterize the spatio-temporal dynamics in the model, we considered peak timing and synchrony in model epidemic results. Peak timing of influenza epidemics across spatial areas is shown to be early in the baseline and travel models, and shifted to a later time in the school closure and holiday models (Figure 5A). The baseline model also highlights spatial variability in peak timing across metro areas, with lower variability in the holiday and school closure models. We then compared the distribution of flu incidence before, during, and after the holiday periods to characterize the trajectory of the model epidemics. The baseline model showed increasing disease incidence from before to after the holiday period, as the baseline epidemic is uninterrupted by the holiday. We also observed that the baseline and travel models appear to show spatial asynchrony, illustrated by the large variance in the distribution of flu incidence across locations, compared with the relatively low variance (spatial synchrony) of the school closure and holiday models (Figure 5B).

**Figure 5:**
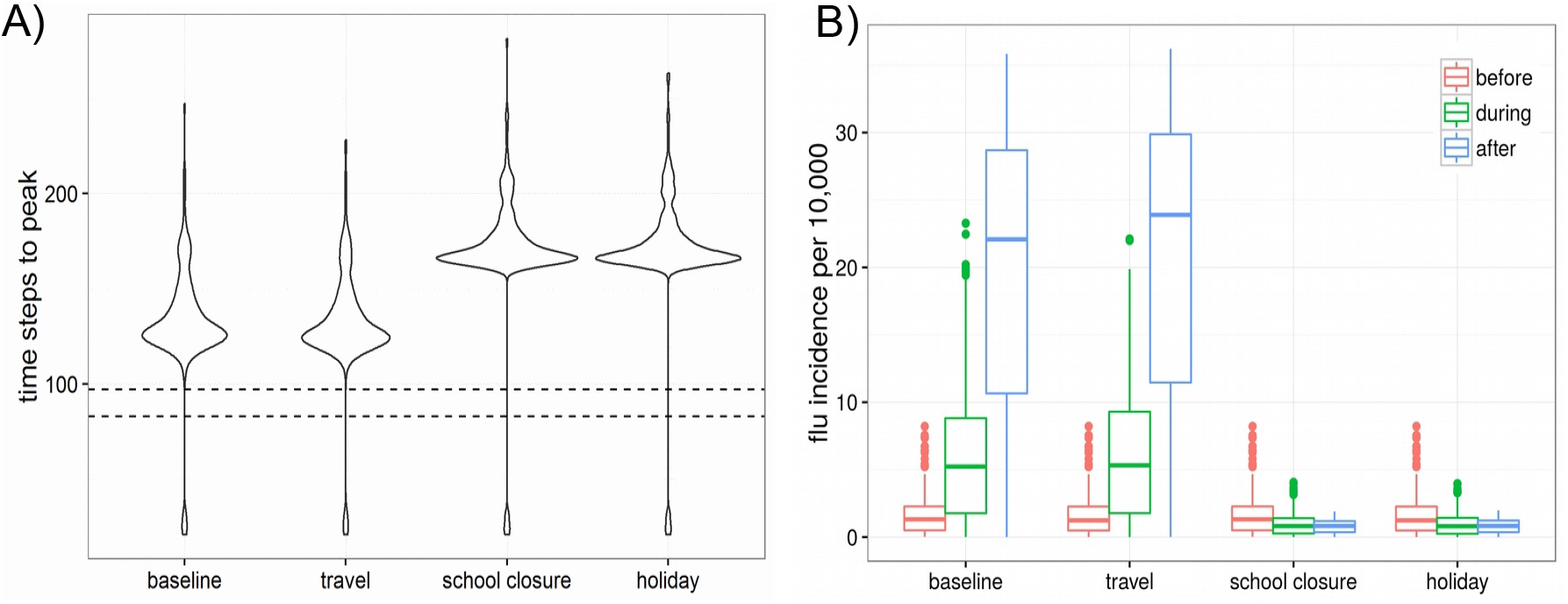
Holiday-associated behavioral changes delay peak timing and increase the synchrony of epidemics across locations in model simulations. **A**) Distribution of time steps (days) to peak across all metro areas, averaged across all model runs. Distributions across metros are compared for the baseline, travel only, contact only, and full holiday models, and the holiday period is demarcated by the horizontal black lines. B) Distributions of flu incidence across all metro areas averaged for the two week durations defined as ‘before’, ‘during’, and ‘after’ the holiday periods, averaged across all model runs. Distributions are compared for the baseline, travel only, school closure only, and full holiday interventions.

To further explore the sensitivity of holiday dynamics to changes in individual-level contact patterns, we considered models in which the overall number of contacts was reduced for children and adults (to match that of the “school closure” model), but the proportion of contact within and between age groups was not altered. We found that this new model (“partial school closure”) produced epidemic dynamics and spatial synchrony patterns matching those of the school closure model (Figure S6). We also explored the sensitivity of our results to the timing of the holiday period, and found that later holidays (occurring during the growth phase or at the epidemic peak) did not impact the overall dynamics, age-specific patterns, or spatial synchrony outcomes reported above (Figures S8, S10, S11), but did result in multiple epidemic peaks due to the resetting of the epidemic trajectory.

## Discussion

In this study, we have considered the impact of winter holiday periods in the United States on influenza transmission. We have observed that these periods are associated with temporary reductions in rates of influenza-like illness, especially among children, at both national and local scales, after controlling for temporal variation in reporting. These observations corroborate previous empirical studies that report reduced flu transmission during the winter holidays in Argentina [17] and in Arizona [18]. Additionally, we observed that the holidays may alter the synchrony of ILI incidence across locations, even for seasons marked by early peaks (occurring before the holiday period).

These empirical patterns are descriptive of the dynamics of influenza during the holiday period, but questions about the mechanisms driving these patterns remain. Our study used an age-specific spatial metapopulation model to compare outcomes with and without temporary, holiday-associated changes in contact and travel. This model framework allowed us to determine which behavioral change drives observed holiday epidemic dynamics and the impact of holiday timing on epidemic outcomes. We found that holiday-associated changes to age-specific contact patterns reproduced many of the patterns observed in our empirical data. In the model, holiday contact patterns were responsible for causing temporary reductions in flu activity during the holiday period, shifting the risk of disease from children to adults, pushing the epidemic peak later in time, and increasing the synchrony in flu incidence during the holiday period. However, total epidemic sizes were not affected (in contrast to the work of [34]). Our analysis also illustrated that the age shift in burden and spatial synchronization caused by holiday periods was insensitive to the timing of the holiday period during the epidemic, but that later holiday periods did result in multiple peaks (as experienced in the 2009 H1N1 pandemic [34].) On the other hand, the empirical patterns from the 2003–2004 early flu season highlight that a holiday period arriving well after an epidemic peak results in minimal impact on epidemic dynamics.

In comparing the effects of two purported mechanisms driving holiday dynamics -school closure and increased travel -we were surprised to find that changes due to school closure explained nearly all of the delay in peak timing and increase in spatial synchrony in our model. Travel restrictions have been shown to produce little to no effect on the spread of pandemic influenza [27, 28, 29], and our study adds to this literature on the minimal effect of holiday-related travel re-routing on spatial influenza transmission. While holiday travel is more commonly linked with seeding and synchronizing flu in multiple locations [24], we found that school closures, and more specifically, reductions in the average number of holiday contacts and not the changes in mixing among age groups, could in fact create a dampening and synchronizing spatial effect (See SM section S3.1). Thus our study lends model-based support to the success of school closures, specifically when they are effectively implemented and timed early during the epidemic [10].

In both empirical and modeling analyses, evidence suggested that children and adults have staggered, temporary dips in reported ILI after the winter holidays; the reduction was later and smaller in magnitude for adults, supporting results that holiday effects are delayed by one week among other age groups relative to children [17], that children are the primary drivers of household transmission [35, 36], and that children experience the greatest disease burden when populations are naive to new strains of influenza and adults are more affected in subsequent seasons or epidemic waves [37, 38]. Nevertheless, the timing of the overall epidemic peak was shifted equally in the holiday model for both children and adults; this suggests that while children may experience greater flu burden and local transmission due to their high contact rates [39, 40, 8, 6], they do not necessarily lead the epidemic wave [41, 5]. Finally, we observed that holiday-associated behavioral changes consistently increased the risk of disease among adults relative to that among children. Our previous work leveraged the consistency of this temporary change to detect early warning signals of seasonal influenza severity [1], and future work could examine how this early flu testbed might signal other actionable epidemiological information about the flu season.

While our holiday model parameters are indeed based on empirical survey data, we have limited knowledge about holiday-associated behavioral changes to contact and travel patterns at a population scale [42, 20, 21, 43]. There are no counterfactual scenarios to observe system behavior in the absence of holidays, but future studies could address this limitation by comparing seasonal flu dynamics across locations with different holiday timings (e.g., different national holidays across countries or different closure timings across school districts). The average holiday travel changes in our model may also not have captured non-routine travel and local travel patterns that have been shown previously to synchronize and spread local epidemics [2, 3, 4], nor did they consider the individual identities of travelers, which additionally slows the speed of epidemic waves [44]. Additionally, flu transmission and ILI reporting were conflated in the model, as the effects of holiday interventions in the model were observed immediately. Future work may account for delays in timing between holiday-associated transmission and reporting (e.g., flu incubation period, time delay before seeking care for ILI) by incorporating these mechanisms into a revised model, or with the use of empirical data at a finer temporal scale. Seasonal environmental fluctuations like temperature and humidity, both of which are hypothesized to modulate influenza transmission and survival, were also not considered among our epidemic impacts [15, 45, 46, 47]. If winter holidays are indeed pushing U.S. epidemics later into the winter as our results demonstrate, environmental conditions may be more favorable to flu transmission. Future work should extend the examination of the link between environmental factors and holiday-induced shifts in peak timing [48, 49, 50]. Additionally, we acknowledge that populations poorly captured by medical claims data (e.g., uninsured individuals) may engage in systematically different holiday-associated behaviors than other populations, as lack of health insurance and financial barriers may be correlated with holiday activities [51, 52]. Future studies with U.S. medical claims may better represent the entire population, as the Affordable Care Act has substantially expanded insurance coverage among adults [53, 54].

## Footnote page

### Supporting Material

Additional Document with detailed descriptions about data processing, model parameters, sensitivity analyses, and supporting evidence.

### Funding

This work was supported by the Jayne Koskinas Ted Giovanis Foundation for Health and Policy (JKTG) [dissertation support grant to ECL]; and the RAPIDD Program of the Science & Technology Directorate, Department of Homeland Security and the Fogarty International Center, National Institutes of Health. The content is solely the responsibility of the authors and does not necessarily reflect the official views of JKTG, the National Institutes of Health, or IMS Health.

## Acknowledgments

The authors thank IMS Health for kindly sharing the medical claims data with us and Vittoria Colizza for her insightful comments on an earlier draft of this work.

## Corresponding author

Shweta Bansal

Assistant Professor, Department of Biology

Georgetown University

; 202-687-9256

## Authors’ contributions

AE parsed the medical claims data, assembled the demographic and contact data, performed analyses, interpreted the results, and wrote the first draft of the manuscript. ECL assembled all other data and performed analyses. CV interpreted the results and edited the manuscript. SB and ECL jointly conceived and designed the study, guided the analysis, interpreted the results, and edited the manuscript. All authors read and approved the final manuscript.

